# Alevin efficiently estimates accurate gene abundances from dscRNA-seq data

**DOI:** 10.1101/335000

**Authors:** Avi Srivastava, Laraib Malik, Tom Smith, Ian Sudbery, Rob Patro

## Abstract

We introduce alevin, a fast end-to-end pipeline to process droplet-based single cell RNA sequencing data, which performs cell barcode detection, read mapping, unique molecular identifier deduplication, gene count estimation, and cell barcode whitelisting. Alevin’s approach to UMI deduplication accounts for both gene-unique reads and reads that multimap between genes. This addresses the inherent bias in existing tools which discard gene-ambiguous reads, and improves the accuracy of gene abundance estimates.

---

There has been a steady increase in the throughput of single-cell RNA-seq (scRNA-seq) experiments, with droplet-based protocols (dscRNA-seq) ^1–3^ facilitating experiments assaying tens of thousands of cells in parallel. The three most widely-used dscRNA-seq protocols: drop-seq ^1^, inDrop ^2^, and 10x-chromium ^3^, use two separate barcodes that require appropriate processing for accurate quantification estimation. First, cellular barcodes (CBs) are used to tag each cell with a unique barcode, which enables pooling of cells for sequencing. Second, identification of PCR duplicates is aided by Unique Molecular Identifiers (UMIs), which tag each unique molecule prior to amplification. Appropriately accounting for the barcode information is therefore crucial for accurate estimation of gene expression. Only a minor fraction of the possible CBs present will ultimately tag a cell, and likewise, only a minor fraction of UMIs will tag unique molecules from the same gene. Thus, in each case, the aim is to identify the barcodes used. Unfortunately, both CBs and UMIs are subject to errors that occur during sequencing and amplification ^1,4^, which makes the accurate deconvolution of this information *in silico* a non-trivial task.

Various methods have been proposed to correctly process dscRNA-seq barcodes in an error-aware manner (“whitelisting”) ^3–6^ and to obtain cell-level gene quantification estimates ^7,8^. Here, we describe an end-to-end quantification pipeline that takes as input sample-demultiplexed FASTQ files and outputs gene-level UMI counts for each cell in the library. We call this unified pipeline alevin, outlined in Figure 1, and it overcomes two main shortcomings of traditional pipelines. First, existing techniques for UMI deduplication discard reads that map to more than one gene. In bulk RNA-seq datasets (with paired-end reads and full-length transcript coverage), the proportion of gene ambiguous reads is generally small (Table S1). Yet, in tagged-end scRNA-seq, this set of gene-ambiguous reads is generally larger, and commonly accounts for ~ 14 – 23% of the input data (Figure 2a, Table S2). This is a result of both the fact that dscRNA-seq protocols, by construction, display a very strong 3’ bias and that these protocols yield effectively single-end reads (only one of the sequenced reads contains sequence from the underlying transcript), resulting in a reduced ability to resolve multimapping using a pair of reads from a longer fragment. Discarding the multimapping reads can strongly bias the gene-level counts predicted by various methods (Figure S1) and lead to under-counting (Figure S2) of UMIs, since the UMI deduplication methods used by these methods do not have a mechanism to deal with such UMIs that map between multiple genes. Second, existing quantification pipelines combine independent processing algorithms and tools for each step, usually communicating results between pipeline stages via intermediate files on disk, which significantly increases the processing time and memory requirements for the complete analysis (Figure S3). We show that alevin makes use of more reads than other pipelines (Figure 2a), that this leads to more accurate quantification of genes (Figure 2b), and that alevin does this 10 times faster and with a lower memory requirement (Figure 2c), when compared to existing best practice pipelines for dscRNA-seq analysis.

**Figure 1:**
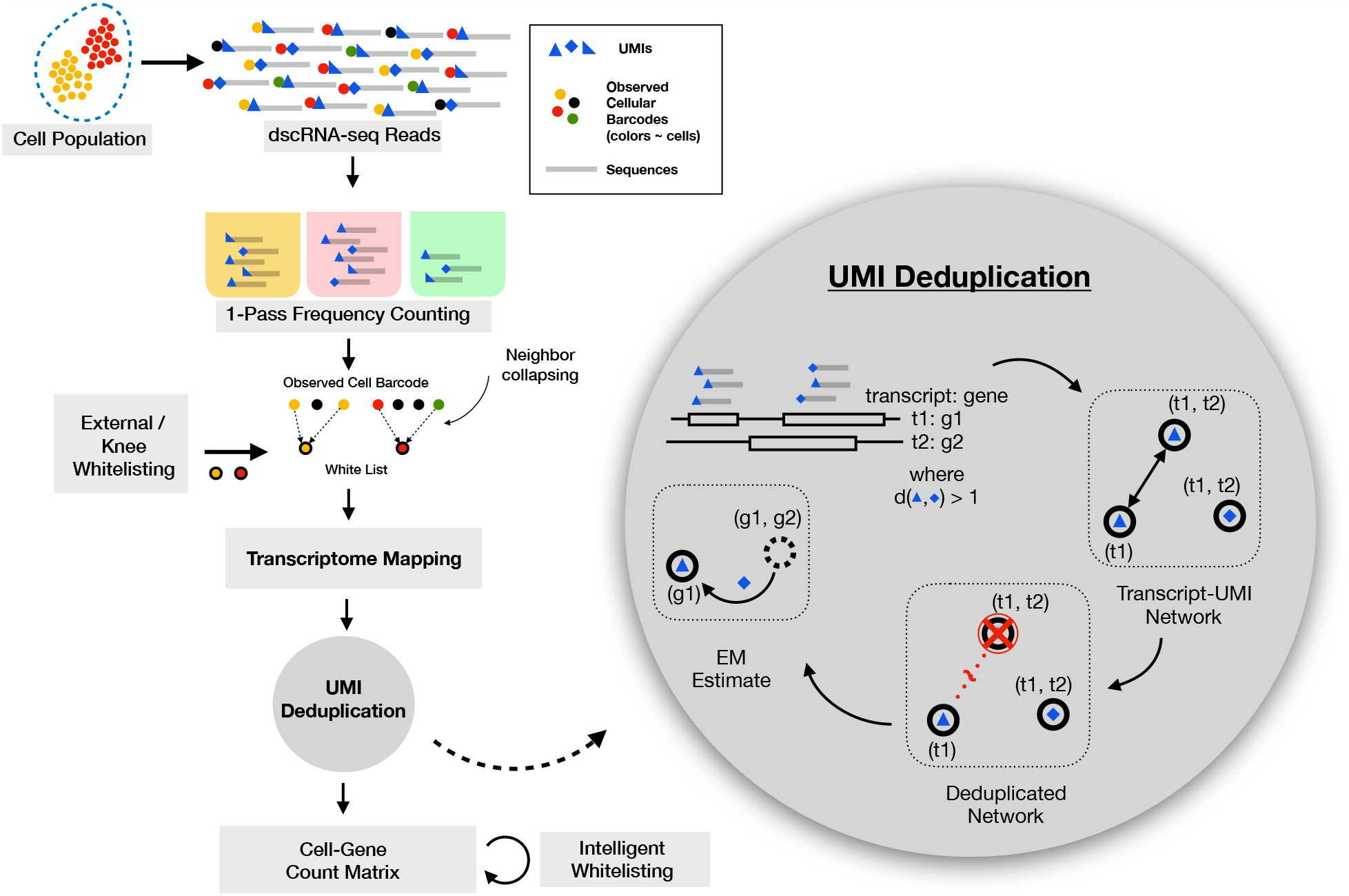
Overview of the alevin pipeline. The input to the pipeline are sample-demultiplexed FASTQ files and there are several steps, outlined here, that are required to process this data and obtain per cell gene-level quantification estimates. The first step is cell-barcode (CB) whitelisting using their frequencies. Barcodes neighboring whitelisted barcodes are then associated with (collapsed into) their whitelisted counterparts. Reads from whitelisted CBs are mapped to the transcriptome and the UMI-transcript equivalence classes are generated. Each equivalence class contains a set of transcripts, the UMIs that are associated with the reads that map to each class and the read count for each UMI. This information is used to construct a graph where each node represents a UMI-transcript equivalence class and nodes are connected based on the associated read counts. The UMI deduplication algorithm then attempts to find a parsimonious set of transcripts that cover the graph (where each consistently-labeled connected component — each monochromatic arborescence) is associated with a distinct pre-PCR molecule. In this way, each node is assigned a transcript label, and in turn, an associated gene label. Reads associated with arborescences that could be consistently labeled by multiple genes are divided amongst these possible loci probabilistically based on an expectation-maximization algorithm. Finally, after obtaining this gene count matrix, an intelligent whitelisting procedure finalizes a list of high quality CBs using a naїve Bayes classifier to differentiate between high and low-quality cells.

**Figure 2:**
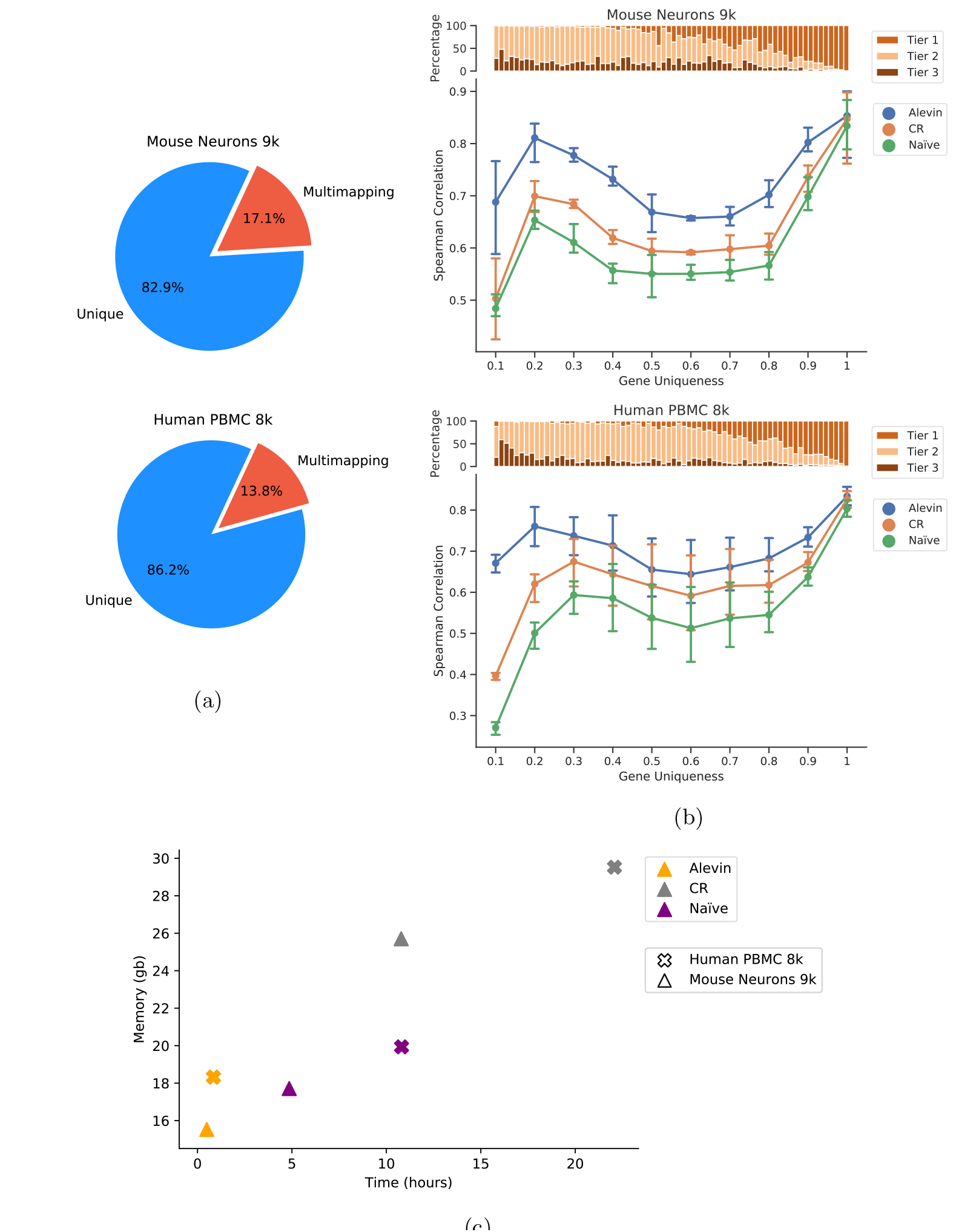
Various analyses on the single cell human PBMC 8k and mouse neuronal 9k datasets. (a) The percentage of multimapping reads in each dataset, calculated using the alignments done by alevin against the transcriptome. A read is considered to multimap if it maps to transcripts from two different genes. (b) The Spearman correlation between quantification estimates (summed across all cells) from different scRNA-seq methods against bulk data from the same cell types, stratified by gene sequence uniqueness. The bar plot shows the percentage of genes in each bin that have unique read evidence. Tier 1 is the set of genes where all the reads are uniquely mapping. Tier 2 is genes that have ambiguously mapping reads, but connected to unique read evidence as well, that can be used by the EM to resolve the multimapping reads. Tier 3 is the genes that have no unique evidence and the read counts are, therefore, distributed between these genes according to an uninformative prior. Note that all methods perform very similarly on genes from tier 1, but the performance of alevin is much better for the other tiers. (c) The time and memory performance of the different pipelines for the two datasets.

There are several steps in the alevin pipeline that are streamlined to work without the overhead of writing to disk, as highlighted in Figure 1 (details in Online Methods). The first step is to identify the CBs that represent properly-captured and tagged cells (“whitelisting”). Alevin uses a two-step whitelisting procedure. An initial whitelist is produced by finding the “knee” in the cumulative distribution of CB frequencies ^1,3^. For each non-whitelisted CB, alevin tries to correct it to a whitelisted CB, otherwise, the barcode and its associated reads are discarded. The next step is read mapping to a target transcriptome ^9,10^, followed by UMI deduplication. Alevin performs deduplication by constructing a graph using information from the UMI sequences, the UMI counts and the transcript equivalence classes ^11^, such that each UMI-transcript equivalence class pair is represented by a node and there exists a directed edge from a node to any node that could have arisen from it due to a PCR or sequencing error, and a bi-directed to any node that could have arisen from it by sampling (without error) from a different position along a duplicate of the same pre-PCR molecule (Section OM2.1). An optimal covering of this graph, using the transcripts associated with each node, will give the minimum number of UMIs, along with their counts, required to explain the set of mapped reads. Since the decision version of this problem is NP-complete (Section S1.6), we propose a greedy algorithm to obtain a minimum cardinality covering of this graph. The ambiguous reads remaining after this UMI resolution phase are assigned based on an expectation-maximization method ^12^. Finally, having obtained per-cell gene expression estimates, CB whitelisting is finalized using a naїve Bayes classifier to differentiate between high and low-quality cells utilizing a set of features derived from the expression estimates and other diagnostic features ^6^(Section OM3). In addition to the gene-by-cell count matrix, alevin also provides information about the reliability of the abundance estimate computed for each gene in each cell in the form of a *tier* matrix (and, optionally, the summarized variance of bootstrap estimates), which succinctly encodes the quality of the evidence used to derive the corresponding count (Section OM2.2).

To assess the performance of alevin, both in terms of accuracy in quantification and resource consumption, we ran it on 10x Chromium datasets from human and mouse, containing 8, 381 and 9, 128 cells and 784 million and 383 million reads respectively. We compare our results against the Cell-Ranger pipeline ^3^ and a custom pipeline, with an external list of whitelisted CBs, using STAR^13^, featureCounts ^14^, and UMI-tools ^4^, which we refer to as the *naïve* pipeline. The exact parameters for running each tool are provided in Section OM4. Results from datasets containing fewer cells are also presented in the Supplementary Material. Comparisons with other pipelines, and on data using the Drop-seq ^1^ protocol, are also detailed in Section S1.7.

To test the accuracy of the quantification estimates, we aggregate the estimates from each of the single-cell quantification tools (summing across all cells) and calculate the correlation with estimates predicted by RSEM ^15^ (paired with Bowite2^16^ alignments) using bulk datasets from the same cell types. Estimates from alevin, when summed across all cells, have a higher Spearman rank correlation than the Cell-Ranger and *naïve* pipelines (Table S6). Specifically, we posit that Cell-Ranger and *naïve* demonstrate a strong and persistent bias against groups of two or more genes that exhibit high sequence similarity. That is, the more sequence-similar a gene is to another gene, the less likely these pipelines are able to assign reads to it — in the extreme case, some genes essentially become *invisible* due to the *in silico* biases of these approaches (a similar effect was reported by Robert and Watson ^17^ in bulk RNA-seq data when simple read-counting approaches are used for quantification, where they highlight that many such genes are relevant to human disease).

To further explore this hypothesis, we stratified the accuracy of the different methods by the uniqueness of the underlying genes (Figure 2b, Figure S6, Section S1.9). In agreement with the hypothesized relationship, we observed that the higher accuracy of alevin is particularly large for genes with a lower proportion of unique k-mers. Thus, the approach of Cell-Ranger and *naïve*, which discard reads mapping to multiple genes, results in systematic inaccuracies in genes which are insufficiently unique (i.e. which share a high degree of sequence homology with some other gene). This bias could impact the expression estimates of important marker genes, such as the genes for the hemoglobin alpha and beta proteins in the mouse neurons ^18,19^. Due to their lower uniqueness ratio, Cell-Ranger appears to exhibit a bias against such genes, and their expression, as predicted by alevin, is systematically higher (Figure S7). Anecdotally, we also noticed that, in the human PBMC data, alevin sometimes predicts the expression of even relatively sequence-unique genes, like YIPF6, that we expect to be expressed in a subpopulation of these cells (monocytes) ^20^, but which exhibit almost no expression as predicted by Cell-Ranger (Figure S8). Because the bias against sequence-ambiguous genes is fundamental and sequence-specific, it cannot be easily remedied with more data, but instead requires the development of fundamentally novel algorithms, like alevin, that account for, rather than discard, reads mapping to such genes. Hence, alevin not only quantifies a greater proportion of the sequenced data than existing methods, but also does so more accurately and in a less-biased manner.

The time and memory requirements for alevin are significantly less than those for the existing pipelines (Figure 2c, Figure S9), where all methods were ran using 16 threads. For the smallest dataset (900 mouse neuronal cells), alevin was ~ 7 times faster than *naïve* and ~ 21 times faster than Cell-Ranger. This difference increases further as the size of the dataset increases, since the performance of alevin scales better than the other tools. Hence, where alevin took only 51 minutes to process the human PBMC 8k dataset, Cell-Ranger took 22 hours and *naï;ve* took 11 hours. In terms of memory, alevin used only ~ 18GB on the human PBMC 8k cell dataset, whereas *naïve* took ~20GB. For the mouse neuronal 9k cell dataset, alevin used 15GB and *naïve* used ~18GB. In both cases, Cell-Ranger required a minimum of 16GB just for STAR indexing. We note that Cell-Ranger allows the user to specify a maximum resident memory limit, and we ran Cell-Ranger allowing it to allocate up to 120GB so that the extra runtime was not due to limitations in available memory. We observe that the optimal number of threads for running alevin is 12 – 16, where the maximum gain in terms of time and memory is achieved (Figure S10).

Our analyses demonstrate that, compared to Cell-Ranger (and *naïve*), alevin achieves a higher accuracy, in part because of considering a substantially larger number of reads. Further, alevin is considerably faster and uses less memory than these other approaches. These speed improvements are due to a combination of the fact that alevin uses bespoke algorithms for CB and UMI edit distance computation, read mapping, and other tasks, and is a unified tool for performing all of the initial processing steps, obviating the need to read and write large intermediate files on disk. Alevin is written in C++14, and is integrated into the salmon tool available at https://github.com/COMBINE-lab/salmon.

## Acknowledgments

The authors would like to thank Fatemeh Almodaresi and Hirak Sarkar for useful discussions during the development of the alevin method, and would also like to thank Hirak Sarkar for his help in crafting Figure 1. This work was supported by the US National Science Foundation (BIO-1564917, CCF-1750472, CNS-1763680), and the US National Institutes of Health (R01HG009937). This project has been made possible in part by grant number 2018-182752 from the Chan Zuckerberg Initiative DAF, an advised fund of Silicon Valley Community Foundation. The authors would like to thank Stony Brook Research Computing and Cyberinfrastructure, and the Institute for Advanced Computational Science at Stony Brook University for access to the high-performance SeaWulf computing system, which was made possible by a $1.4M National Science Foundation grant (#1531492).

## Online Methods

### OM1 Initial whitelisting and barcode correction

After standard quality control procedures, the first step of existing single-cell RNA-seq processing pipelines ^1;2;3^ is to extract cell barcode and UMI sequences, and to add this information to the header of the sequenced read or save it in temporary files. This approach, while versatile, can create many intermediate files on disk for further processing, which can be time and space-consuming.

Alevin begins with sample-demultiplexed FASTQ files. It quickly iterates over the file containing the barcode reads, and tallies the frequency of all observed barcodes (regardless of putative errors). We denote the collection of all observed barcodes as *B*. Whitelisting involves determining which of these barcodes may have derived from a valid cell. When the data has been previously processed by another pipeline, a whitelist may already be available for alevin to use. When a whitelist is not available, alevin uses a two-step procedure for calculating one. An initial draft whitelist is produced using the procedure explained below, to select CBs for initial quantification. This list is refined after per-cell level quantification estimates are available (see section OM3) to produce a final whitelist.

To generate a putative whitelist, we follow the approach taken by other dscRNA-seq pipelines by analyzing the cumulative distribution of barcode frequencies, and finding the knee in this curve ^1;2^. Those barcodes occurring after the knee constitute the whitelist, denoted 𝒲. We use a Gaussian kernel to estimate the probability density function for the barcode frequency and select the local minimum corresponding to the “knee”. In the case of a user-provided whitelist, the provided 𝒲 is used as the fixed final whitelist.

Next, we consider those barcodes in *ℰ* =*B*\ 𝒲 to determine, for each non-whitelisted barcode, whether a) its corresponding reads should be assigned to some barcode in 𝒲 or b) this barcode represents some other type of noise or error (e.g., ambient RNA, lysed cell, etc.) and its associated reads should be discarded. The approach of alevin is to determine, for each barcode *h_j_* ∈ *ℰ*, the set of whitelisted barcodes with which *h_j_* could be associated. We call these the putative labels of *h_j_* — denoted as *l*(*h_j_*). Following the criteria used by previous pipelines ^1^, we consider a whitelisted barcode *w_i_* to be a putative label for some erroneous barcode *h_j_* if *h_j_* can be obtained from *w_i_* by a substitution, by a single insertion (and clipping of the terminal base) or by a single deletion (and the addition of a valid nucleotide to the end of *h_j_*). Rather than applying traditional algorithms for computing the all-versus-all edit-distances directly, and then filtering for such occurrences, we exploit the fact that barcodes are relatively short. Therefore, we can explicitly iterate over all of the valid *w_i_* ∈ *𝒲* and enumerate all erroneous barcodes for which this might be a putative label. Let *Q*(*w_i_,H*) be the set of barcodes from *ℰ* that adhere to the conditions defined above; then, for each *h_j_* ∈ *Q*(*w_i_,H*), we append *w_i_* as putative label for the erroneous barcode *h_j_*.

Once all whitelisted barcodes have been processed, each element in *ℰ* will have zero or more putative labels. If an erroneous barcode has more than one putative label, we prioritize substitutions over insertions and deletions. If this does not yield a single label, ties are broken randomly. If no candidate is discovered for an erroneous barcode, then this barcode is considered “noise”, and its associated reads are simply discarded.

### OM2 Mapping reads and UMI de-duplication

After labeling each barcode, either as noise, or as belonging to some whitelisted barcode, alevin maps the sequenced reads to the target transcriptome ^4;5^. Reads mapping to a given transcript (or multimapping to a set of transcripts) are categorized hierarchically, first based on the label of their corresponding cellular barcode, and then based on their unique molecular identifier (UMI). At this point, it is then possible to deduplicate reads based on their mapping and UMI information.

The process of read deduplication involves the identification of duplicate reads based on their UMIs and alignment positions. Most amplification occurs prior to fragmentation in library construction for 10x Chromium protocols ^6^. Because of this, the alignment position of a given read is not straightforward to interpret with respect to deduplication, as the same initial unique molecule may yield reads with different alignment coordinates^†^. UMIs can also contain sequence errors. Thus, achieving the correct deduplication requires proper consideration of the available positional information and possible errors.

Our approach for handling sequencing errors, and PCR errors in the UMIs is motivated by “directional” approach introduced in UMI-tools ^7^. Let *𝒰 _i_* be the set of UMIs observed for gene *i*. A specific UMI *u_n_* ∈ *𝒰 _i_*, observed *c_n_* times in gene *i*, is considered to have arisen by PCR or sequence error if there exists *u_m_* ∈ *𝒰 _i_* such that *d*(*u_n_, u_m_*) = 1 and *c_m_* > 2*c_n_* + 1, where *d*(*·,·*) is, the Hamming distance. Using this information, only UMIs that could not have arisen as an error under this model are retained. However, this approach may over-collapse UMIs if there exists evidence that similar UMIs (i.e. UMIs at a Hamming distance of 1 or less) may have arisen from different transcripts, and hence, distinct molecules. Moreover, this approach first discards reads that multimap to more than one read, causing it to lose a substantial amount of information before even beginning the UMI deduplication process.

As previously proposed to address the problem of cell-clustering ^8^, an equivalence class ^9;10;11;12;13;14;15^ encodes some positional information, by means of encoding the set of transcripts to which a fragment is mapped. Specifically, these equivalence classes can encode constraints about which UMIs may have arisen from the same molecule and which UMIs — even if mapping to the same gene — must have derived from distinct pre-PCR molecules. This can be used to avoid over-collapsing UMI tags that are likely to result from different molecules by considering UMIs as distinct for each equivalence class. However, in its simplest form, this deduplication method is prone to reporting a considerably higher number of distinct UMIs than likely exist (Figure S5). This is because reads from different positions along a single transcript, and tagged with the same UMI, can give rise to different equivalence classes, so that membership in a different equivalence class is not, alone, sufficient evidence that a read must have derived from a distinct (pre-PCR) molecule (Section S1.7.2). This deters us from directly using such a UMI collapsing strategy for deriving gene-level counts, though it may be helpful for other types of analyses.

Given the shortcomings of both approaches to UMI deduplication, we propose, instead, a novel UMI resolution algorithm that takes into account transcript-level evidence when it exists, while simultaneously avoiding the problem of under-collapsing that can occur if equivalence classes are treated independently for the purposes of UMI deduplication.

#### OM2.1 UMI Resolution Algorithm

A potential drawback of gene-level deduplication is that it discards transcript-level evidence. In this case, such evidence is encoded in the equivalence classes. Thus, gene-level deduplication provides a conservative approach and assumes that it is highly unlikely for molecules that are distinct transcripts of the same gene to be tagged with a similar UMI (within an edit distance of 1 from another UMI from the same gene). However, entirely discarding transcript-level information will mask true UMI collisions to some degree, even when there is direct evidence that similar UMIs must have arisen from distinct transcripts. For example, if similar UMIs appear in transcript-disjoint equivalence classes (even if all of the transcripts labeling both classes belong to the same gene), then they *cannot* have arisen from the same pre-PCR molecule. Accounting for such cases is especially true when using an error-aware deduplication approach, and as sequencing depth increases.

To perform UMI deduplication, alevin begins by constructing a graph *G* = (*V*, *E*), for each cell, where each *v_i_* = (*u, T_i_*) is a tuple consisting of UMI sequence *u* and a set of transcripts 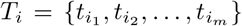. There is a count associated with each vertex such that *c*(*v_i_*) = *c_i_* is the number of times this UMI, equivalence class pair is observed. *G* contains two types of edges; directed and bi-directed. There exists a directed edge between every pair of vertices (*v_i_, v_j_*) for which *c_i_* > 2*c_j_ -* 1, |*T_i_* ∩ *T_j_*| > 0, and *d*(umi(*v_i_*), umi(*v_j_*)) = 1. There is a bi-directed edge between every pair of vertices (*v_k_*, *v_l_*) for which *d*(umi(*v_k_*), umi(*v_l_*)) = 0 and |*T_k_* ∩ *T_l_*|> 0. Once the edges of this graph have been formed, we no longer need to consider the counts of the individual UMI, equivalence class pairs.

We view the problem of determining the true number of distinct molecules appearing in this cell, as well as the mapping relationship between UMIs and genes, as follows. Motivated by the principle of parsimony, we wish to explain the observed vertices (i.e., UMI, equivalence class pairs) via the minimum possible number of pre-PCR molecules that are consistent with the observed data. We pose this problem in the following manner. Given a graph *G*, we seek a *minimum cardinality covering by monochromatic arborescences*. In other words, we wish to cover *G* by a collection of vertex-disjoint arborescences (the analog of a rooted tree in a directed graph), where each arborescence is labeled consistently by a set of transcripts, which are the pre-PCR molecule types from which its reads and UMIs are posited to have arisen. Further, we wish to cover all vertices in *G* using the minimum possible number of arborescences. Here, the graph *G* defines which UMI, read pairs can potentially be explained in terms of others (i.e. which vertices may have arisen from the same molecule by virtue of different fragmentation positions or which vertices may have given rise to other through PCR duplication with error). The decision version of this problem is NP-complete (Section S1.6), and so alevin employs a greedy algorithm in practice to obtain a valid, though not necessarily minimum, covering of *G*.

The algorithm employed by alevin works as follows. First, we note that weakly-connected components of *G* can be processed independently, and so we describe here the procedure used to resolve UMIs within a single weakly-connected component — this is repeated for all such components. Let *C* = (*V_C_*, *E_C_*) denote our current component. We perform a breadth-first search starting from each vertex *v_i_* ∈ *V_C_* and considering each transcript 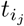 (the *j*^th^ transcript in the equivalence class labeling vertex *v_i_*). We compute the size (cardinality) of the largest arborescence that can be created starting from this node and using this label to cover the visited vertices. Let 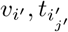 be the vertex, transcript pair generating the largest arborescence and let 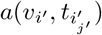 be the corresponding arborescence. We now remove all of the vertices in 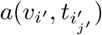, and all of their incident edges, from *C*, and we repeat the same procedure on the remaining graph. This process is iterated until all vertices of *C* have been removed. This procedure is guaranteed to select some positive order arborescence (i.e. an arborescence containing at least one node) in each iteration, and hence is guaranteed to terminate after at most a linear number of iterations in the order of *C*.

After computing a covering, each arborescence is labeled with a particular transcript. However, the selected transcript may not be the unique transcript capable of producing this particular arborescence starting from the chosen root note. We can compute, for each arborescence, the *set* of possible transcript labels that could have colored it (i.e. those in the intersection of the equivalence class labels for all of the vertices in the arborescence). If the cardinality of this set is 1, then only a single transcript is capable of explaining all of the UMIs associated with this arborescence. If the cardinality of this set is > 1, then we need to determine if all transcripts capable of covering this arborescence belong to the same gene, or whether transcripts from multiple genes may, in fact, be capable of explaining the associated UMIs. In the former case, the count of pre-PCR molecules (i.e. distinct, deduplicated UMIs) associated with this uniquely-selected gene is incremented by 1. In the latter case, the molecule associated with the arborescence is considered to potentially arise from any of the genes with which it could be labeled. Subsequently, an EM algorithm is used to distribute the counts between the genes. Note that other pipelines simply discard these gene-ambiguous reads and that both manners in which alevin attempts to resolve such reads (i.e. either by being selected via the parsimony condition or probabilistic allocated by the EM algorithm) are novel in the context of scRNA-seq quantification. The EM procedure we adopt to resolve ambiguous arborescences proceeds in the same manner as the EM algorithm used for transcript estimation in bulk RNA-seq data ^13^, with the exception that we assume the probability of generating a fragment is directly proportional to the estimated abundance, rather than the abundance divided by the effective length (i.e. we assume that, in the tagged-end protocols used, there is no length effect in the fragment generation process).

#### OM2.2 Tier Assignment

The alevin program also outputs a tier matrix, of the same dimensions as the cell-gene count matrix. Within a cell, each gene is assigned one of fours tiers. The first tier (assigned 0) is the set of genes that have no read evidence in this cell and are, therefore, predicted to be unexpressed (whether truly absent, or the effect of some dropout process). The rest of the tiers (1,2, and 3) are assigned based on a graph induced by the transcript equivalence classes as follows:

1. All equivalence classes of size 1 are filtered out. The genes associated with the transcripts from these classes are assigned to tier 1.
2. For the remaining equivalence classes, of size > 1 gene, a graph *G* is constructed. The nodes in *G* are transcripts and two nodes share an edge if their corresponding transcripts belong to a single equivalence class.
3. All the connected components in *G* are listed and the transcript labels on the nodes mapped to their corresponding genes. If any component contains a node whose gene has previously been assigned to tier 1, that gene and all other genes in this connected component are assigned to tier 2. Hence, tier 2 contains genes whose quantification is impacted by the EM algorithm (after the UMI deduplication).
4. Genes associated with the remaining nodes in the graph are assigned tier 3. These are genes that have no unique evidence, and do not share reads (or, in fact, paths in the equivalence class graph) with another gene that has unique evidence. Hence, the EM algorithm will distribute reads between these genes in an essentially uniform manner, and their estimates are uninformative. Their abundance signifies that some genes (at least 1) in this ambiguous family are expressed, but exactly which and their distribution of abundances cannot be determined.

Alevin, optionally (using the --numCellBootstraps flag), also outputs bootstrap variance estimates for genes within each cell. These variance estimates could conceivably be used by downstream tools for dimensionality reduction, differential expression testing, or other tasks.

### OM3 Final whitelisting

Many existing tools for whitelisting CBs, such as Cell-Ranger ^3^ and Sircel ^16^ perform whitelisting only once. As discussed above, both tools rely on the assumption that the number of times a CB is observed is sufficient to identify the *correct* CBs, i.e. those originating from droplets containing a cell. However, as observed by Petukhov et al.^17^, there is considerable variation in sequencing depth per-cell, and some droplets may contain damaged or low-quality cells. Thus, true CBs may fall below a simple knee-like threshold. Similarly, erroneous CBs may lie above the threshold. Petukhov et. al ^17^ proposed that instead of selecting a single threshold, one should treat whitelisting as a classification problem and segregate CBs into three regions; high-quality, low-quality and uncertain / ambiguous. Here, high-quality refers to the CBs which are deemed to be definitely correct, and low-quality are the CBs which are deemed to most likely not arise from valid cells. A classifier can then be trained on the high and low-quality CBs to classify the barcodes in the ambiguous region as either high or low-quality. We adopt this approach in alevin, using our knee method’s cutoff to determine the ambiguous region. Specifically, we divide everything above the knee threshold into two equal regions; high-quality valid barcodes (upper-half), denoted by *ℋ*, and ambiguous barcodes (lower-half), denoted by *ℒ* To learn the low-quality region, we take *n_l_* = max(0.2 · |*ℋ*|, 1000) barcodes below the knee threshold.

In the implementation of Petukhov et al.^17^, a kernel density estimation classifier was trained using features which described the number of reads per UMI, UMIs per gene, the fraction of intergenic reads, non-aligned reads, the fraction of lowly expressed genes and the fraction of UMIs on lowly expressed genes. In addition, a maximum allowable mitochondrial read content was set for a CB to be classified as “high-quality”. Whilst these features enabled the authors to build a classifier which efficiently separated “high-quality” cells from “low-quality” cells, we believe it may be possible to improve this set of features. Specifically, most of these features would be expected to correlate with the number of reads or UMIs per CB. Thus, the classifier is biased towards attributes associated with higher read depth, when in fact one wants it to learn the feature attributes associated with high-quality cells. We therefore used a slightly different set of features which we believe may better capture the differences between high and low-quality cells:

1. Fraction of reads mapped
2. Fraction of mitochondrial reads
3. Fraction of rRNA reads
4. Duplication rate
5. Mean gene counts post de-duplication
6. The maximum correlation of gene-level quantification estimates with the high-quality CBs. (Optionally activated by `--useCorrelation flag`.)

We chose to use a naїve Bayes classifier to perform classification, since we observed no clear difference between multiple ML methods (not shown), and the naїve Bayes classifier yields classification probabilities which are easy to interpret. Our final set of whitelisted CBs are those classified as high-confidence. We observe that the number of high-confidence cells predicted by alevin are in close proximity to the count of cells predicted by Cell-Ranger, but there are non-trivial differences (Table S8).

### OM4 Machine Configuration and Pipeline Replicability

10x v1 chemistry benchmarking has been scripted using Snakemake ^18^ and performed on an Intel(R) Xeon(R) CPU (E5-2699 v4 @2.20GHz with 44 cores and 56MB L3 cache) with 512GB RAM and a 4TB TOSHIBA MG03ACA4 ATA HDD running Ubuntu 16.10.

10x v2 chemistry benchmarking has been scripted using CGATCore (https://github.com/cgat-developers/cgat-core). The full pipeline and analysis are performed using Stony Brook’s seawulf cluster with 164 Intel Xeon E5-2683v3 CPUs.

For all analyses, the genome and gtf versions used for human datasets was GENCODE release 27, GRCh38.p10 and for mouse datasets was GENCODE release M16, GRCm38.p5. All transcriptome files were generated using these with “rsem-prepare-reference”.

All relevant details of running the following single cell tools are available at https://github.com/COMBINE-lab/alevin-paper-pipeline/tree/master/benchmark_pipeline.

#### Cell-Ranger (v2.2.0)

The following additional flags were used, as recommended by the Cell-Ranger guidelines: `--nosecondary --expect-cells NumCells`, where NumCells is 10,000 for PBMC 8k and Neurons 9k, 5,000 for PBMC 4k, 2,000 for Neurons 2k and Neurons 900.

#### Alevin (v0.12.0)

Run with default parameters with the chromium protocol flag and the -lISR to specify strandedness. The mRNA and rRNA lists were obtained from the relevant annotation files and passed as input. Experiments on v1 chemistry were run using the same flags but with the `--gemcode protocol flag`.

#### STAR (v2.6.0a)

The following flag was used, as recommended by the guidelines of UMI-tools: `--outFilterMultimapNmax 1`

#### featureCounts (v1.6.3)

This was run to obtain an output BAM file and with stranded input (-s 1).

#### UMI-tools (v0.5.4)

The extract command was used to get the CBs/UMIs, when provided with an external CB whitelist, and attach it to the corresponding reads. The following flags were used in the count command to obtain the per cell gene count matrix: `--gene-tag=XT --wide-format-cell-counts`

#### Dropseq utils (v2.0.0)

All the commands were run as recommended by the authors in the tool’s manual.

The bulk datasets were quantified using Bowtie2 and RSEM, run as follows:

#### Bowtie2 (v2.3.4.3)

The following flags were used, as recommended in the guidelines of RSEM: `--sensitive --dpad 0 --gbar 99999999 --mp 1,1 --np 1 --score-min L,0,-0.1 --no-mixed --no-discordant`

#### RSEM (v1.3.1)

Run with default parameters.

## Footnotes

† We note that whether the majority of amplification occurs pre- or post-fragmentation can be protocol specific and can suggest different strategies for UMI deduplication. Here, we are primarily concerned with the 10X Chromium protocols, dominated by pre-fragmentation amplification. However, the method we propose for UMI deduplication can be applied to other protocols as well.

## References

[1] Evan Z Macosko, Anindita Basu, Rahul Satija, James Nemesh, Karthik Shekhar, Melissa Goldman, Itay Tirosh, Allison R Bialas, Nolan Kamitaki, Emily M Martersteck, et al. Highly parallel genome-wide expression profiling of individual cells using nanoliter droplets. Cell, 161(5):1202–1214, 2015.

[2] Allon M Klein, Linas Mazutis, Ilke Akartuna, Naren Tallapragada, Adrian Veres, Victor Li, Leonid Peshkin, David A Weitz, and Marc W Kirschner. Droplet barcoding for single-cell transcriptomics applied to embryonic stem cells. Cell, 161(5):1187–1201, 2015.

[3] Grace XY Zheng, Jessica M Terry, Phillip Belgrader, Paul Ryvkin, Zachary W Bent, Ryan Wilson, Solongo B Ziraldo, Tobias D Wheeler, Geoff P McDermott, Junjie Zhu, et al. Massively parallel digital transcriptional profiling of single cells. Nature Communications, 8:14049, 2017.

[4] Tom Smith, Andreas Heger, and Ian Sudbery. UMI-tools: modeling sequencing errors in unique molecular identifiers to improve quantification accuracy. Genome Research, 27(3):491–499, 2017.

[5] Lu Zhao, Zhimin Liu, Sasha F Levy, and Song Wu. Bartender: a fast and accurate clustering algorithm to count barcode reads. Bioinformatics, 2017.

[6] Petukhov, V and Guo, J and Baryawno, N and Severe, N and Scadden, DT and Samsonova, MG and Kharchenko, PV. dropEst: pipeline for accurate estimation of molecular counts in droplet-based single-cell RNA-seq experiments. Genome Biology, 19(1):78, 2018.

[7] Vasilis Ntranos, Govinda M Kamath, Jesse M Zhang, Lior Pachter, and N Tse David. Fast and accurate single-cell RNA-seq analysis by clustering of transcript-compatibility counts. Genome Biology, 17(1): 112, 2016.

[8] Luyi Tian, Shian Su, Xueyi Dong, Daniela Amann-Zalcenstein, Christine Biben, Azadeh Seidi, Douglas J Hilton, Shalin H Naik, and Matthew E Ritchie. scPipe: a flexible R/Bioconductor preprocessing pipeline for single-cell RNA-sequencing data. PLoS Computational Biology, 14(8):e1006361, 2018.

[9] Avi Srivastava, Hirak Sarkar, Nitish Gupta, and Rob Patro. RapMap: a rapid, sensitive and accurate tool for mapping RNA-seq reads to transcriptomes. Bioinformatics, 32(12):i192–i200, 2016.

[10] Hirak Sarkar, Mohsen Zakeri, Laraib Malik, and Rob Patro. Towards selective-alignment: Bridging the accuracy gap between alignment-based and alignment-free transcript quantification. In Proceedings of the 2018 ACM International Conference on Bioinformatics, Computational Biology, and Health Informatics, BCB ’18, pages 27–36, New York, NY, USA, 2018. ACM. ISBN 978-1-4503-5794-4. doi: 10.1145/3233547.3233589. URL http://doi.acm.org/10.1145/3233547.3233589.

[11] Ernest Turro, Shu-Yi Su, Ângela Gonçalves Lachlan JM Coin, Sylvia Richardson, and Alex Lewin. Haplotype and isoform specific expression estimation using multi-mapping RNA-seq reads. Genome Biology, 12(2):R13, 2011.

[12] Rob Patro, Geet Duggal, Michael I Love, Rafael A Irizarry, and Carl Kingsford. Salmon provides fast and bias-aware quantification of transcript expression. Nature Methods, 14(4):417, 2017.

[13] Alexander Dobin, Carrie A Davis, Felix Schlesinger, Jorg Drenkow, Chris Zaleski, Sonali Jha, Philippe Batut, Mark Chaisson, and Thomas R Gingeras. STAR: ultrafast universal RNA-seq aligner. Bioinformatics, 29(1):15–21, 2013.

[14] Yang Liao, Gordon K Smyth, and Wei Shi. featureCounts: an efficient general purpose program for assigning sequence reads to genomic features. Bioinformatics, 30(7):923–930, 2013.

[15] Bo Li and Colin N Dewey. RSEM: accurate transcript quantification from RNA-Seq data with or without a reference genome. BMC Bioinformatics, 12(1):323, 2011.

[16] Ben Langmead and Steven L Salzberg. Fast gapped-read alignment with bowtie 2. Nature Methods, 9 (4):357, 2012.

[17] Christelle Robert and Mick Watson. Errors in RNA-Seq quantification affect genes of relevance to human disease. Genome Biology, 16(1):177, 2015.

[18] Xiaoping Han, Renying Wang, Yincong Zhou, Lijiang Fei, Huiyu Sun, Shujing Lai, Assieh Saadatpour, Zimin Zhou, Haide Chen, Fang Ye, et al. Mapping the mouse cell atlas by microwell-seq. Cell, 172(5): 1091–1107, 2018.

[19] Franziska Richter, Bernhard H Meurers, Chunni Zhu, Vera P Medvedeva, and Marie-Françoise Chesselet. Neurons express hemoglobin *α* and *β*-chains in rat and human brains. Journal of Comparative Neurology, 515(5):538–547, 2009.

[20] Helder I Nakaya, Jens Wrammert, Eva K Lee, Luigi Racioppi, Stephanie Marie-Kunze, W Nicholas Haining, Anthony R Means, Sudhir P Kasturi, Nooruddin Khan, Gui-Mei Li, et al. Systems biology of vaccination for seasonal influenza in humans. Nature Immunology, 12(8):786, 2011.

## References

[4] Avi Srivastava, Hirak Sarkar, Nitish Gupta, and Rob Patro. RapMap: a rapid, sensitive and accurate tool for mapping RNA-seq reads to transcriptomes. Bioinformatics, 32(12):i192–i200, 2016.

[5] Hirak Sarkar, Mohsen Zakeri, Laraib Malik, and Rob Patro. Towards selective-alignment: Bridging the accuracy gap between alignment-based and alignment-free transcript quantification. In Proceedings of the 2018 ACM International Conference on Bioinformatics, Computational Biology, and Health Informatics, BCB ’18, pages 27–36, New York, NY, USA, 2018. ACM. ISBN 978-1-4503-5794-4. doi: 10.1145/3233547.3233589. URL http://doi.acm.org/10.1145/3233547.3233589.

[6] 10x-genomics single-cell 3’-v2 kit. https://teichlab.github.io/scg_lib_structs/data/CG000108_AssayConfiguration_SC3v2.pdf.

[7] Tom Smith, Andreas Heger, and Ian Sudbery. UMI-tools: modeling sequencing errors in unique molecular identifiers to improve quantification accuracy. Genome Research, 27(3):491–499, 2017.

[8] Vasilis Ntranos, Govinda M Kamath, Jesse M Zhang, Lior Pachter, and N Tse David. Fast and accurate single-cell RNA-seq analysis by clustering of transcript-compatibility counts. Genome Biology, 17(1): 112, 2016.

[9] Ernest Turro, Shu-Yi Su, Ângela Gonçalves, Lachlan JM Coin, Sylvia Richardson, and Alex Lewin. Haplotype and isoform specific expression estimation using multi-mapping RNA-seq reads. Genome Biology, 12(2):R13, 2011.

[10] Aziz M Mezlini, Eric JM Smith, Marc Fiume, Orion Buske, Gleb L Savich, Sohrab Shah, Sam Aparicio, Derek Y Chiang, Anna Goldenberg, and Michael Brudno. iReckon: simultaneous isoform discovery and abundance estimation from RNA-seq data. Genome Research, 23(3):519–529, 2013.

[11] Rob Patro, Stephen M Mount, and Carl Kingsford. Sailfish enables alignment-free isoform quantification from RNA-seq reads using lightweight algorithms. Nature Biotechnology, 32(5):462, 2014.

[12] Nicolas L Bray, Harold Pimentel. Páll Melsted, and Lior Pachter. Near-optimal probabilistic RNA-seq quantification. Nature Biotechnology, 34(5):525, 2016.

[13] Rob Patro, Geet Duggal, Michael I Love, Rafael A Irizarry, and Carl Kingsford. Salmon provides fast and bias-aware quantification of transcript expression. Nature Methods, 14(4):417, 2017.

[14] Zhaojun Zhang and Wei Wang. RNA-Skim: a rapid method for RNA-Seq quantification at transcript level. Bioinformatics, 30(12):i283–i292, 2014.

[15] Chelsea J-T Ju, Ruirui Li, Zhengliang Wu, Jyun-Yu Jiang, Zhao Yang, and Wei Wang. Fleximer: Accurate quantification of RNA-Seq via variable-length k-mers. In Proceedings of the 8th ACM International Conference on Bioinformatics, Computational Biology, and Health Informatics, pages 263–272. ACM, 2017.

[16] Akshay Tambe and Lior Pachter. Barcode identification for single cell genomics. BioRxiv, page 136242, 2017.

[17] Petukhov V and Guo J and Baryawno N and Severe N and Scadden DT and Samsonova MG and Kharchenko PV. dropEst: pipeline for accurate estimation of molecular counts in droplet-based single-cell RNA-seq experiments. Genome Biology, 19(1):78, 2018.

[18] Johannes Köster and Sven Rahmann. Snakemake—a scalable bioinformatics workflow engine. Bioinformatics, 28(19):2520–2522, 2012.

